# Community Detection in Brain Connectome using Quantum Annealer Devices

**DOI:** 10.1101/2022.12.21.521454

**Authors:** Marcin Wierzbiński, Joan Falcó-Roget, Alessandro Crimi

## Abstract

Recent advancements in network neuroscience are pointing in the direction of considering the brain as a small-world system with segregated regions integrated to facilitate different cognitive tasks and functions. In this context, community detection is a pivotal issue in computational neuroscience. In this paper we explore community detection within brain connectomes using the power of quantum annealers, and in particular the Leap’s Hybrid Solver. Our results shows that quantum annealers can achieve higher modularity index compared to classical annealer while computing communities of brain connectomes. Those promising preliminary results points out that quantum annealers might be the better choice compared to classical computing optimization process.

## 1 Introduction

Network neuroscience is an emerging discipline that tries to examine brain organizing principles using network science tools. It was made possible by the intersection of networks science and neuroscience^1^. This allows the merging of two worlds which have shown tremendous advancements in recent years. The first world is given by complex system studies through graph analysis^2^, the second is given by neuroimaging, which permits modeling of brain structure and function even representing relevant information as graph^3^. Community (also called cluster or module) detection is an explored yet not solved task in many fields using graph representations^4^. Previous research has demonstrated how a modular structure can be used to highlight links between topological and functional aspects of complex networks that are not trivial^5^. The human brain is a complex network made up of physically connected areas that can be activated when performing certain tasks or while at rest. Indeed, in network neuroscience, the most used connectivity representations are the *functional* and *structural* representations, though other representations exist, as the effective connectivity which attempts to combine these two. More specifically, *functional connectivity* represents the covarying activity of spatially separated brain areas observed in time series data as functional magnetic resonance imaging or electroencephalogram^6^. A graph is then built defining nodes as brain regions and edges according to the correlation of brain activity. *Structural connectivity* is referred to the physical pathways bridging brain areas, typically evaluated in-vivo by diffusion-weighted imaging^7^, or ex-vivo with staining and histological imaging^8^. In this case, a graph is defined as nodes also given by the brain regions and edges are the physical neuronal pathways. *Effective connectivity* refers to the impact one neurological system has over another, either at the level of individual neurons or entire brain regions and incorporates structural and functional information as it combines the neuronal activities related by physical pathways^9^. We do not focus particularly in this last type of connectivity as many recent criticisms raised concerns about its validity, as it can be seen as just temporal correlation^10^. The most plausible structural organization describing the complexity of brain functions is emerging as the integration of segregated regions (communities)^11^. Therefore, detecting communities in all types of connectivity is relevant, even though what community means in neuroscience is still an open question lacking absolute ground-truth or based on specific characteristic gathering group of neurons^5^.

Previous research in this context has focused on representing properties of isolated brain networks focusing on network features important to brain function. Those include variants of Newman’s modularity function^2^, and its maximization through Louvain-like algorithms^12^ for the detection of clusters of regions^13^. More advanced approaches have expanded this considering brain networks as dynamic and not static networks^14^, and information flow at different time scales^15^. Indeed defining communities in absolute terms has been discouraged in many fields^16^ especially considering possible nodes which can overlap communities and mislabeling. Truly, the brain is not composed by strictly segregated regions with defined regions as Broadmann introduced in 1909^17^, but by a more intricate and integrate complex of segregated networks with smooth borders^18^.

The present work is not focused on defining the ideal community detection algorithm in brain connectomes, but rather we investigate if better clustering results can be achieved using quantum devices. This is motivated by the fact that modularity maximization heuristics have a demanding optimization problem at their core. Moreover, optimal solutions have been shown to be elusive to classical computing devices^19^.

Optimization problems in quantum devices rely on minimization and sampling from energy-based models via combinatorial techniques. The 2-communities modularity maximization problem is closely related to an Ising-like model^20,21^. Furthermore, its generalization to a *k*-state Potts model is exploitable for Simulated Annealing^22^ as well as its quantum partner, namely Quantum Annealing (QA) algorithms. Consequently, we start by formulating the relationship between modularity and energy functions to later exploit them for detecting multiple communities in structural and functional brain graphs. Considering the Louvain algorithm as a reliable state-of-the-art estimate of the true result, we compare its performance in a desktop computer with results obtained with the D-Wave quantum computer^20^. More specifically, we tested on a real quantum annealer device, and not on simulators. Lastly, we consider networks with deterministic connections on quantum devices, and not quantum inspired networks on classical computing^23,24^.

## 2 Methods

In this section we first revise in detail how modularity is defined and how it is mapped into energy based model for both quantum and classical computing (Subsection 2.1 and 2.2). Then we clarify how our community detection is implemented inside the quantum annealer (Subsection 2.23).

Let us define a brain connectome as *G* = (*V, E*) a weighted graph with nodes *i, j* ∈ *V* and edges (*i, j*) ∈ *E*. With *V* being brain regions defined by an atlas, and *E* edges a set of either physical neuronal pathways or correlation of brain activity, such that the corresponding adjacency matrix **A** is defined as follows:

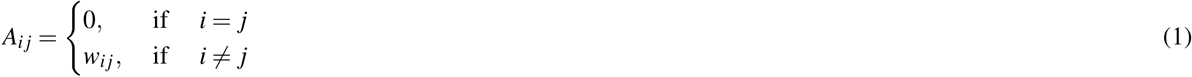

with *w*_*i j*_ representing the weight for edge between nodes *i* and *j*. Those weights are the representation of number of neuronal fiber bundles or intensity of activity correlation. In our case, we define *w*_*i j*_ ∈ *{*0, 1*}* as a binary variable taking into account the existence of a connection between two nodes *i* and *j*. Moreover, the degree of a node is defined as *g*_*i*_ = ∑ _*j*_ *A*_*i j*_, and it measures the number of connections for the node *i* quantified by summing over all its neighbors.

### 2.1 Modularity

Suppose that a graph contains |*V* | = *n* vertices and is divided into *k* communities or clusters denoted as *{C*_1_, …, *C*_*k*_*}*. Given a specified partition, the modularity is defined as^22,25^

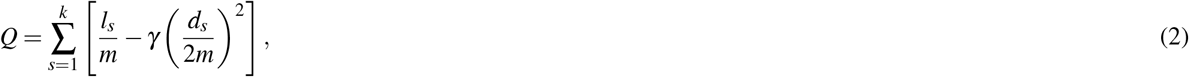

where 2*m* = ∑_*i*_ *g*_*i*_ is the total number of edges,

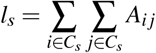

is the number of links between nodes inside cluster *C*_*s*_ and

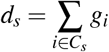

is the sum of the degrees of the nodes belonging to the cluster *C*_*s*_. The resolution parameter *γ* defines the arbitrary trade-off between intra- and inter-group edges^26^. Conveniently^27^, and without loss of generality it can be set to 1, although we comment on other cases in later sections. The modularity as expressed in Eq. (2) is very convenient if the communities present in the graph are already known. However, if the number of communities *k* is unknown, the summation over *s* becomes a problem. To bypass this limitation, one can define a binary variable identifying nodes and/or links that are part of the same cluster. The most widely used proposal was defined in^2^.

For the sake of clarity, we start by having only two communities *{C*_1_,*C*_2_*}*. In this simple case, let *s*_*i*_ = 1 if vertex *i* belongs to community *C*_1_ and *s*_*i*_ = −1 if it belongs to community *C*_2_. We can now define a variable 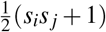 that returns 1 if and only if vertices *i, j* belong to the same community and 0 otherwise. Thus, the total number of existing edges between nodes in the same community can be rewritten as

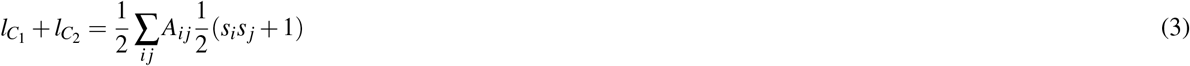

where the 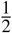 factor has been added to avoid double counting. Moreover, the sum of the degrees in all communities can be easily replaced using the same procedure

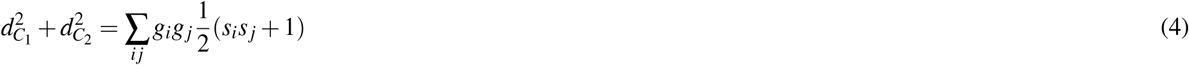

where the double summation over *i j* has been used instead of (∑_*i*∈*s*_ *g*_*i*_)^2^ for completeness. With these changes we can express the modularity *Q* in Eq. (2) as

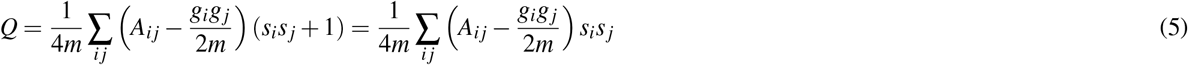

where the rightmost equality follows from 2*m* = ∑_*i*_ *g*_*i*_ = ∑_*i j*_ *A*_*i j*_.

Despite replacing a single summation for two, Eq. (5) has two main advantages, the first one being its easy interpretability. In a graph, if edges are placed at random, the expected number of links between two nodes *i* and *j* can be shown to be precisely 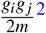. Hence, the modularity *Q* can be understood as the difference between the actual number of edges and those expected by chance inside all the different communities. Second, proposing different sets of two communities is now straightforward. According to Eq. (5), modifying the variable *s*_*i*_ of even a single node yields a completely different partition and corresponding modularity. We will explore the latter in more detail in the following section.

Furthermore, in many graphs, and especially in brain networks, the number of communities is expected to be considerably higher than just a couple. A straightforward generalization can be achieved by replacing the binary variable in the case of just two communities 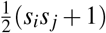 for a community agnostic Kronecker delta function *δ* (*c*_*i*_, *c* _*j*_), then the modularity is expressed as

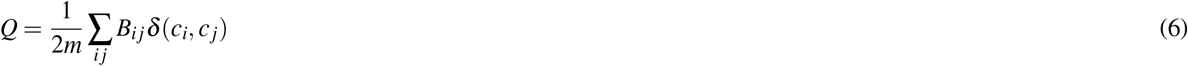

where **B** is known as the modularity matrix whose elements are given by

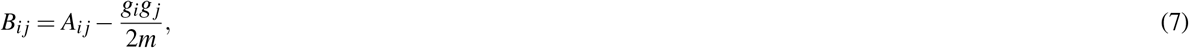

and

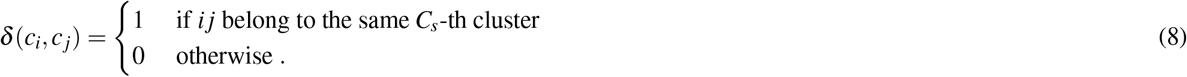

Eventually, the partition in *k* communities is chosen by the maximum modularity score *Q* in function of *k*. Note that permutation of the rows and columns of an adjacency matrix **A** does not alter the graph nor its modularity. However, the output of an algorithm whose goal is to find the partition with highest modularity, could depend on how this adjacency matrix was defined initially. Interestingly, results on many test cases in literature show that the ordering of the nodes does not have a significant influence therefore showing an almost perfect deterministic behavior^12,25^.

### 2.2 Mapping to Energy-Based models

The conundrum of community detection aims at finding the graph partition that maximizes the modularity 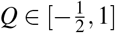. The problem can be solved using brute force by simply trying out all possible combinations of the variables *s*_*i*_, *s* _*j*_ (or ***δ***(*c*_*i*_, *c*_*j*_) for more than two communities), computing the associated modularities and eventually choosing the highest one. However, even for rather small graphs, this approach becomes unfeasible due to a rapid growth in complexity.

Alternative algorithms, such as the ones we use in the current work, use heuristic reasoning^12^ or exploration of state and energy spaces^21,22^. To use the latter, one needs to map the modularity function to a physical system. The simplest of such systems is known as an Ising model, shown to be very useful to study properties of phase transitions in magnetic systems. Consider a system with *N* particles whose magnetic spin is characterised by a discrete variable *s*_*i*_ ∈ {−1, 1}. The energy of the given system immersed in an external magnetic field **h** is given by

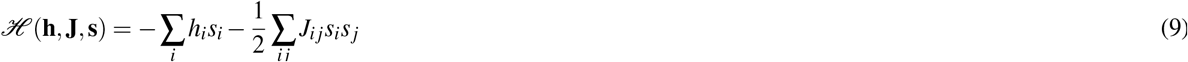

where *s*_*i*_ are the spins of each particle, *h*_*i*_ are the local magnetic fields and *J*_*i j*_ encodes the couplings strengths that encode the interactions between particles *i* and *j*. Usually, Eq. (9) is defined in a regular lattice only considering interactions between 1-st neighbours. Note, nonetheless, that the formulation is agnostic towards the specific spatial topology of the system. Taking this into consideration, it is direct to see that the modularity in Eq. (5) corresponds to an Ising system defined in a non-regular graph, with no external field and coupling matrix 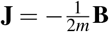.

Furthermore, an Ising model can be easily extended to include more than two states. In this case, each *i*-th particle is fully characterised by a discrete variable *s*_*i*_ ∈ {0,…, *k* − 1} where *k* ∈ ℤ^+^ is a positive integer denoting the number of possible states.

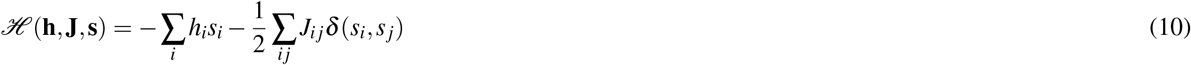

where *h*_*i*_ is the local magnetic field and the Kronecker function is 1 when two particles are in the same state (i.e., *s*_*i*_ = *s* _*j*_). Once again, the modularity in Eq. (6) straightforwardly matches a physical system known as Potts model with the same coupling constant 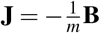 and no external field. A physical system is said to be in thermodynamic equilibrium if the energy of the configuration, given by the successive spin variables *s*_*i*_, rests at a minimum. The problem of how to find this *ground state* has been widely studied and solved using different methods including, but not limited to, QA. In this subsection, we have reviewed how computing the modularity of a certain network organization is equivalent to obtaining the energy of a given magnetic system. Thereafter, we use the same procedures and strategies to find partitions that maximized the modularity or, alternatively, minimized its opposite in Eqs. (5) and (6). That is, we explored configurations *C*_*s*_ such that the corresponding modularity *Q*\*satisfied:

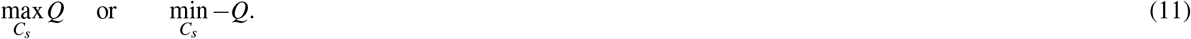

### 2.3 *k* -community detection with a QA processor

QA^28^ relies on the preparation of a physical quantum system whose Hamiltonian evolves according to

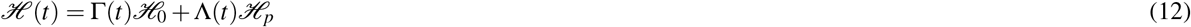

where the ground state of ℋ_0_ is “easy” to prepare and to measure experimentally. The functions Γ and Λ are both continuous and monotonously decreasing and increasing respectively such that ℋ(0) = ℋ_0_ and ℋ(*T*) = ℋ_*p*_. If initially the system is prepared in a way that lies at the ground state of ℋ_0_, the process in Eq. (12) is adiabatic and [ℋ_0_, ℋ_*p*_] ≠ 0 (i.e., the Hamiltonians do not commute), given the adiabatic theorem of quantum mechanics, the system should remain in the ground state at all times, thus ending up in the minimum of ℋ*p*. In this state, any measurement of the system’s elements shall return this ground state configuration. We provide a simplified description of this process in Fig. 1 (a).

**Figure 1.**
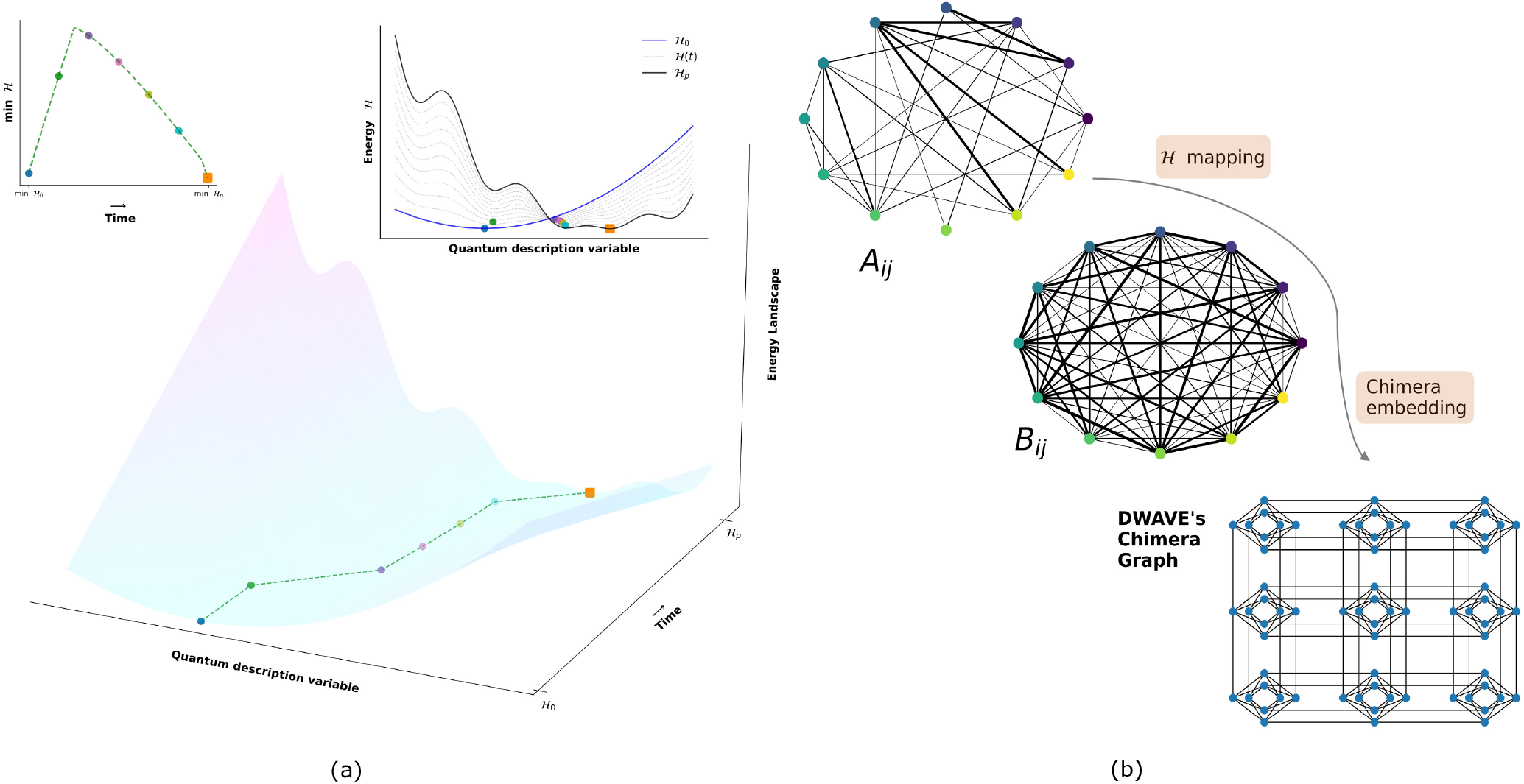
Simplified description of a Quantum Annealing (QA) process in (a). Initially, the system is prepared in the ground truth of an “easy” hamiltonian ℋ_0_ (solid blue line), in this case a simple cuadratic function of a variable describing the quantum state. The hamiltonian of the system, as in Eq (12), is let to evolve adiabatically (dashed grey lines) resting at the minimum of each intermediate hamiltonian ℋ(*t*) (coloured circles). Under these conditions, the final state, depicted as the big orange square, will correspond to the ground state of the desired system ℋ_*p*_ (solid black line). Note that the two discrete jumps (green-purple and cyan-orange) are possible thanks to quantum tunneling. In (b), schematic representation of the logical process explained in the Methods. To partition a graph via QA, the connections of the original network, given by an adjaceny matrix as in Eq. (1), need to be mapped to a fully connected Ising-like system of spins. Note that the dimensions of the Ising system are the same as the original network (identical node coloring) but the strengths of the connections (solid weighted black lines) are now given by the modularity matrix in Eq. (7). The system is then embedded in a *Chimera* graph^20^ (i.e., the quantum processor) by using the DWAVE platform. Each node in the Chimera graph represents a qubit.

The D-Wave hardware, known as a *Chimera* graph, consists of *i* = 1, …, *N* qubits coupled with weights **J** whose energetic state ℋ _*p*_ is described with an Ising Hamiltonian^20^. Therefore, any optimization problem to be solved using quantum annealing necessitates the reduction (or embedding) from its original description (e.g., (6)) to an enclosed embedded form of Eq. (9) in the *Chimera* graph (i.e., a mapping from *G* to ℋ_*p*_). The DWAVE platform^1^ autonomously embeds the DQM version of the problem to the processor (see Fig. 1 (b)).

The modularity maximization problem naturally maps to a Potts model, also known as a Discrete Quadratic Model (DQM). However, a DQM variable *s*_*i*_ takes 1 of *k* possible discrete positive values which is not exactly an Ising spin. Yet, an alternative representation using one-hot encodings facilitates the previously mentioned reduction. A vector **x**_*i*_ is defined such that *x*_*iu*_ = 1 if *s*_*i*_ = *u* and 0 otherwise. This way we obtain a quadratic model subject to the constraint ∑_*u*_ *x*_*iu*_ = 1 that is easily transformed to an Ising representation with the mapping *s*_*iu*_ = 2*x*_*iu*_ − 1. Below, we provide snips of the Python code that reduces the original network object to an Ising model with which the D-Wave system performs the optimization to better understand how it is implemented. Fortunately, the D-Wave platform provides a straightforward way to do this. We used Networkx^29^ to compute the modularity matrix from an adjacency matrix **A** in Eq. (1). The reduction is based on the modularity matrix **B** in Eq. (7) and the expression of the coupling matrix **J** for the quadratic term in Eq. (10). Recall that the external field **h** need not be added. The *DiscreteQuadraticModel* class contains this model, and its methods provide convenient utilities with the specific representation of a problem.

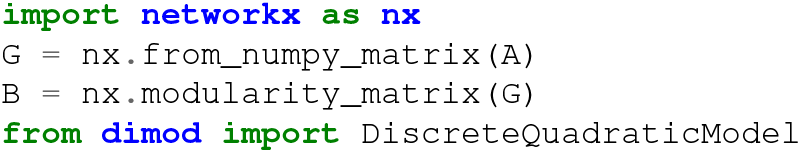
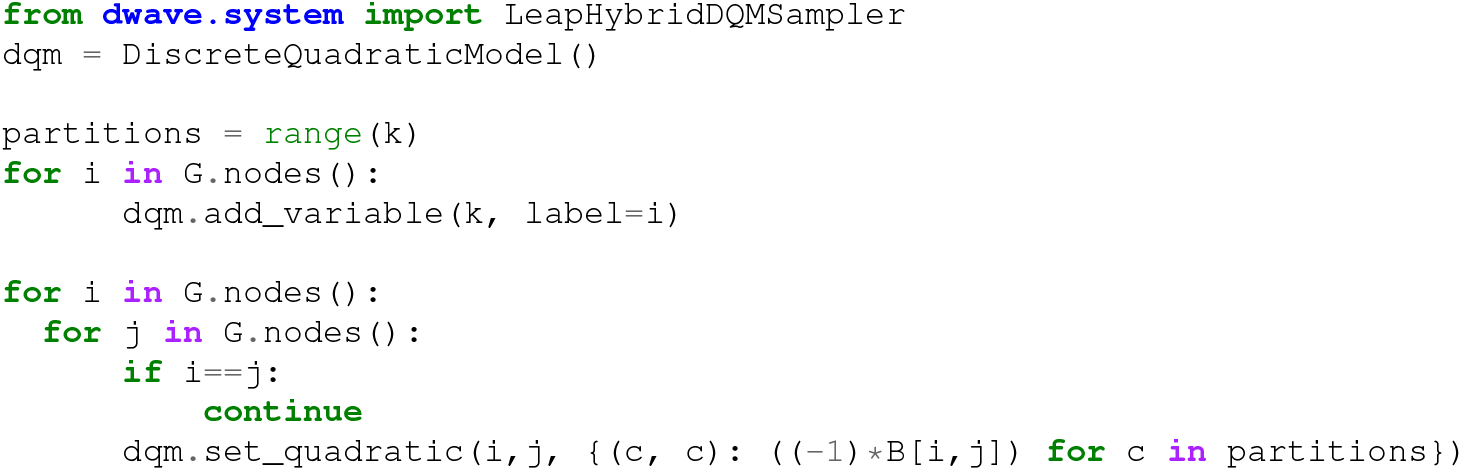

The nested loop is used to express all the possible communities a single node can be assigned to as well as to set the corresponding coupling weight given by *B*_*i j*_. Note that the total number of edges *m* is constant and thus it does not modify the relevant ground state. The code, also skips self-referenced nodes. The *token* can be obtained by logging into https://cloud.dwavesys.com/leap/. The final result is a sample set assigning each node to a particular community.

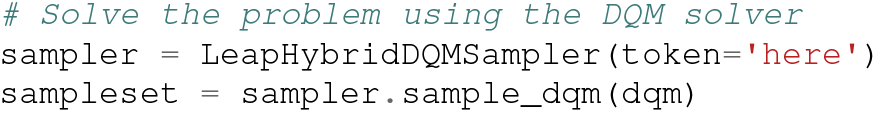

We have used the Leap Hybrid Solver developed by D-Wave. This Solver service can read and solve much larger inputs than the current-model D-Wave 2000Q. This approach leveraged the unique problem-solving capabilities of the Quantum Process Unit and extended those capabilities to larger and more varied types of inputs than would otherwise be possible. It can be called directly through the D-Wave Ocean application programming interface (API). The resulting strings of discrete variables are translated based on the optimization problem’s representation.

In our experiments we investigated how similar the detected communities were to an external proxy both using classical and quantum computing. Where no ground-truth is available, we considered the results of the eigengap, from spectral communities analysis, to be the aforementioned proxy. With the eigengap heuristic, the number of communities is usually given by the value of *k* that maximizes the difference between two consecutive eigenvalues. We kept track of the three largest eigengaps. Then, we computed the modularity index obtained by 100 runs. The third eigengap acted as a limit to see whether the optimization identified the same number of communities or reached higher modularities with less communities.

Finally, we run the same calculation using a Louvain Community Detection Algorithm (LCDA) to solve the community detection problem. The Louvain Community Detection Algorithm is a state-of-the-art heuristic to extract the community structure of a network based on modularity optimization^12^. Given the nature of one of the dataset described later as representing a debated ground-truth^16^, we consider both the case of known and unknown number of communities for this dataset. As in previous similar studies^30,31^, we consider higher modularities as a proxy for better results.

### 2.4 Data and Code availability

In this study, we performed experiments using 2 datasets. The first is a well-known dataset (Zachary karate club^32^) which has been investigated in several community detection studies (e.g.^25^). The second dataset is given by the brain connectomes comprised in the BrainNet Viewer Toolbox defined from atlases^33^, which are the main focus of the study. Both datasets are based on previously acquired data from previous studies, and our analysis represents a numerical simulation study for which no ethical approval was required. The karate club dataset was created by Zachary by observing 34 members of a karate club over a period of 2 years, where a partition of the members happened, and a network of friendships between members of the club was observed. The dataset has been traditionally defined as a ground-truth subdivision of 2 communities^32^ as shown in Supplementary Figure S1. However, communities defined by a specific ground-truth labeling can be misleading and not necessarily reflect the network topology^16^, and in fact higher number of communities in this dataset has been investigated^34^. Therefore, we consider both the cases whether this labeling should be restricted to 2 communities and not.

This toy example is really useful to validate the modularity in case of a known information flow example. The dataset is freely available at the URL https://networkrepository.com/soc-karate.php. The brain connectivity data are obtained from the examples of BrainNet Viewer available at the URL https://www.nitrc.org/projects/bnv/. More specifically, the Automated Anatomical Labeling (AAL) which is the most common atlas for structural connectivity^35^ comprising 90 regions of interest and the Dosenbach atlas which is a functional atlas with 160 regions of interest. For the details of how those regions are used as nodes to construct the networks within BrainNet Viewer, we refer to their paper^33^. There is no particular reason to use this dataset apart the fact that those are brain connectomes publicly available. The basic properties of the networks used in the experiments are reported in Supplementary Table S1.

The code was written in Python version 3.9.7, and it is available at the repository https://github.com/alecrimi/clustering-dwave, it comprises a series of custom scripts and the usage of the Networkx library version 2.8.3^36^, and the Dwave-system library version 1.10.

The used hardware were respectively for quantum and classical computing: the D-Wave 2000Q with Chimera edges, and a powerful workstation with 11th Gen Intel(R) Core(TM) i9-11900KF @ 3.50GHz processor.

## 3 Results

In this section, we report the results obtained by using the aforementioned tools on the 2 datasets. We report here first the resulting eigengap for both datasets, the communities are defined by modularity and the value of modularity.

### 3.1 Karate club dataset

Computing the eigengap Fig. 3 (a) within the karate club dataset we found that the highest gaps were related to *k* = 3, 4, 2. The highest eigengap was given by *k* = 4, while the original defined number of communities is *k* = 2^32^.

**Figure 3.**
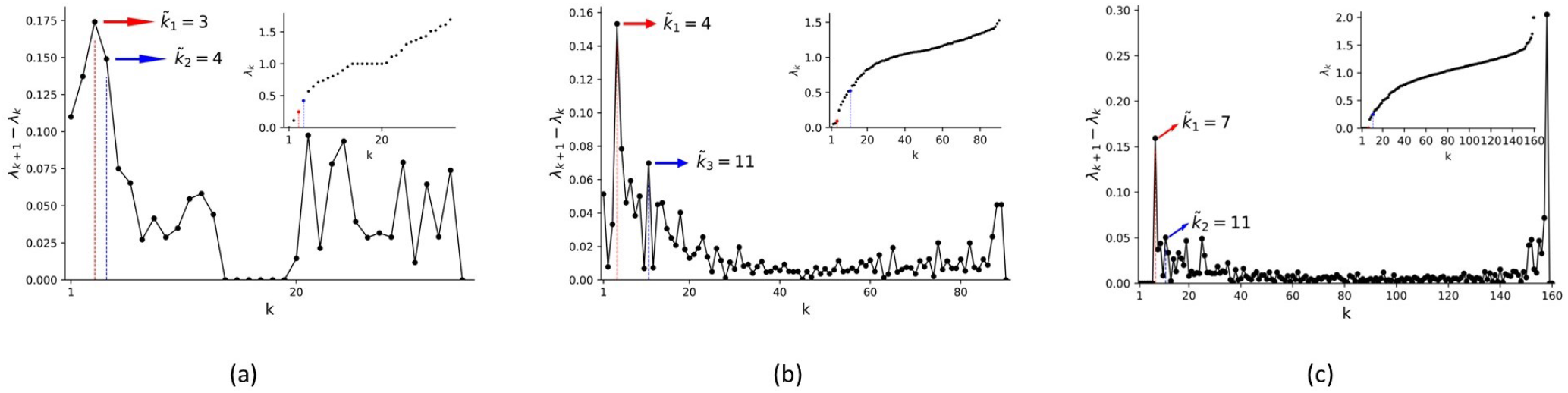
Eigengaps for the Karate club cluster (a), AAL90 brain atlas (b), and Dosenbach brain atlas (c). The red lines highlight the highest gap for each case, while the blue line the second highest gap. For the AAL (in b), the second eigengap is directly visible thus we explicitely mark the third. Last eigenvalues are not taken into consideration.

Forcing both the classical and quantum community detection algorithm to look for 2 communities, both algorithms were able to assign correctly all members of the defined communitiesm (see Fig. S1). It is relevant to mention that to reach this goal, the resolution parameter *γ* in equation (2) had to be set smaller than 1. For all other experiments, we considered the more standard value *γ* = 1.

Following the experiments in an unsupervised manner as described in the previous section, the classical and quantum algorithm showed different results. In Fig. 2 we show the differences of the communities obtained by using classical and quantum computing. It can be seen that only 2 communities (depicted in red and blue) have different grouping. In Table 1 we can see summarized the modularity index for both classical and quantum device. As shown in literature^12,25^, the methods behave in a deterministic matter and as our results show the quantum hardware noise were close to zero even considering a large number of repeated experiments. Therefore, we did not carried out statistical analsys to show the significance of the results reported in Table 1.

**Table 1.** Community detection on karate club dataset. Here, the number of detected communities for classical and quantum computing is the same but the modularity is higher for the quantum annealer used.

| Mod. LCDA | Mean Mod. QA ± std | Max $k$ | $k$ LCDA | $k$ QA |
| --- | --- | --- | --- | --- |
| 0.427 | $0.444 \pm 8.85e-17$ | $k = 4$ | 4 | 4 |

**Figure 2.**
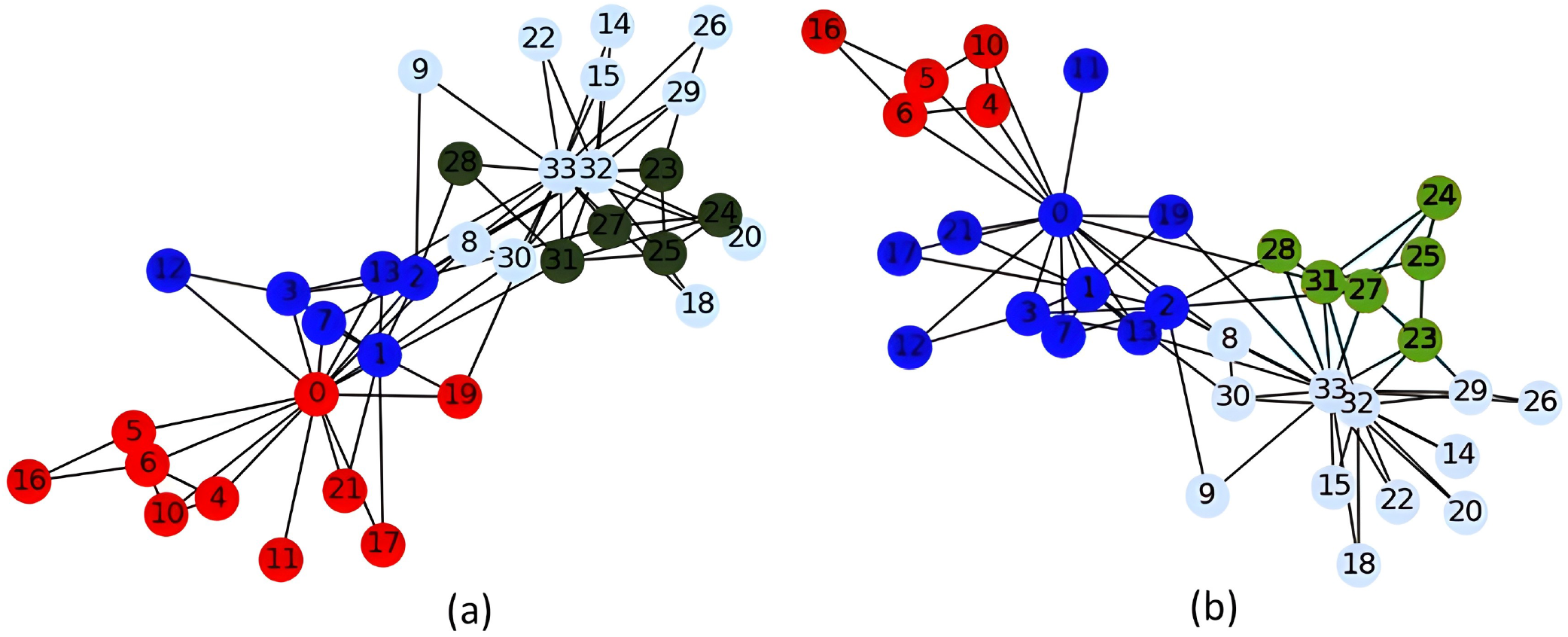
Community detection for the Zachary karate club graph using Louvain Community Detection Algorithm (a), and using Leap Hybrid Solver (b). We can observe that the assignment to communities differs for these 2 methods in the red and blue communities.

### 3.2 Brain connectivity

The eigengaps for the two brain connectome are depicted in Fig. 3 (b) and 3 (c). We found some high eigengaps at the end of the spectrum, namely where we consider each node as a cluster. This might make sense as nodes in brain atlases represent an already relatively large region of interest according to functional activation of cytoarchitecture. Namely, those are already clusters of neurons or brain activity. However, this will not be interesting given the research question of this paper. The 3 largest eigengaps were respectively for the AAL and Dosenbach atlas: 4,5,11 and 7,11,27. Comparing the modularity obtained on the classical and quantum devices with both AAL90 and Dosembach brain connectivity graphs, we found different modularity indexes reported in Table 2. The results were consistently deterministic and separated that a statistical significance analysis was not carried out. The resulting communities are shown in Figs. 4 and 5 respectively for the AAL and Dosenbach atlas. The computational time was for the classical and quantum annealer for the runs related to those networks respectively 0.08 *±* 0.001 and 5.3 *±* 0.08 seconds.

**Table 2.** The results of *k* community detection on benchmark atlases. We show results from both the D-Wave quantum annealer (QA) and a Louvain Community Detection Algorithm (LCDA). The mean and standard deviation (std) for modularity were counted on running the method (QA) 100 times. The LCDA method has the standard deviation zero. This is, the LCDA method returns the same modularity for each run.

| Atlas | Mod. index (LCDA) | Mean Mod. index (QA) ± std | Max clusters $k$ | Optimal $k$ (LCDA) | Optimal $k$ (QA) |
| --- | --- | --- | --- | --- | --- |
| AAL90 | 0.646 | $0.648 \pm 7.85e-17$ | $k = 11$ | 6 | 5 |
| Dosenbach | 0.408 | $0.416 \pm 8.54e-05$ | $k = 27$ | 13 | 9 |

**Figure 4.**
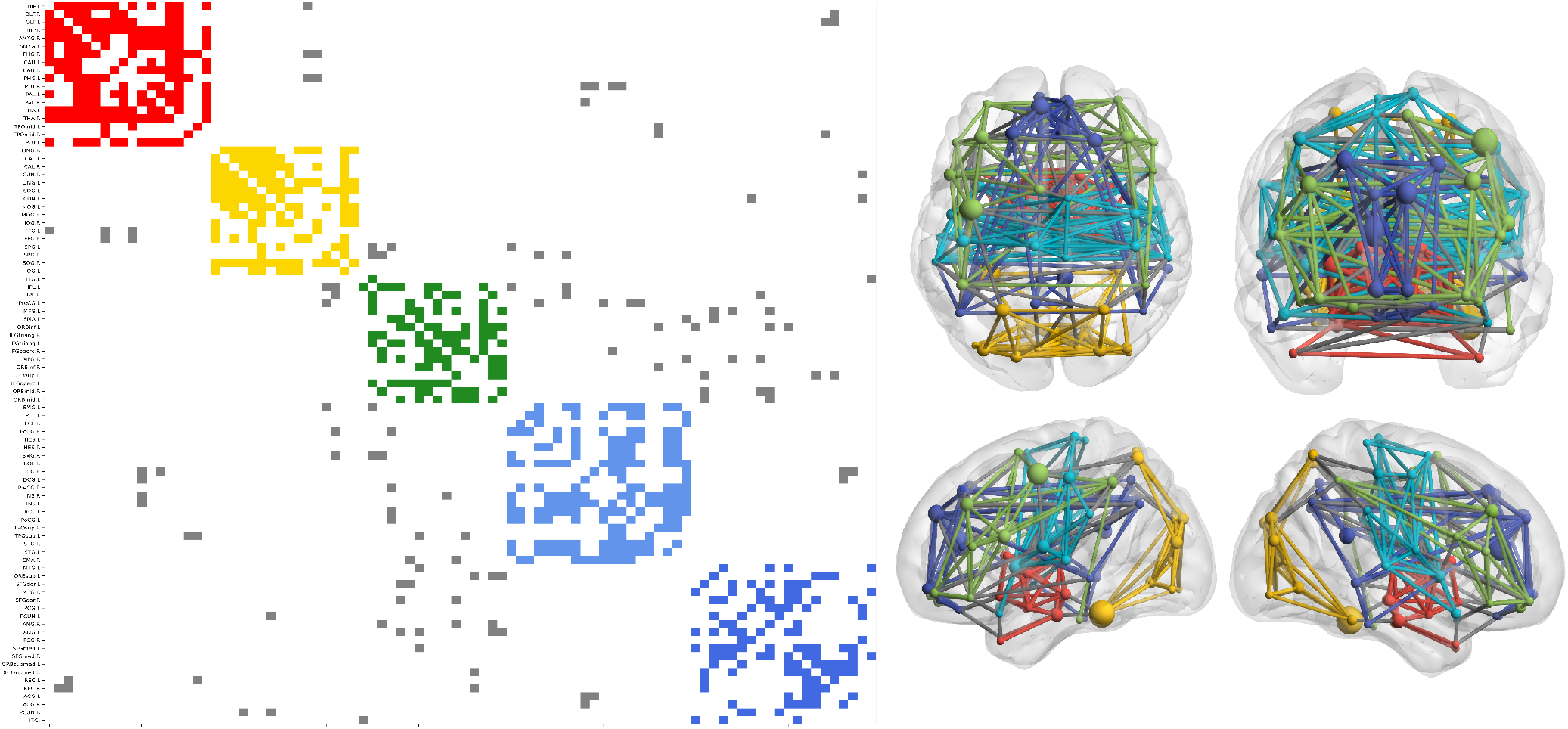
Graph partitioning of the communities defined by using QA for the AAL atlas. Left: Reordered connectivity matrix according to module assignment. Right: Axial, coronal, and 2 hemispheres sagittal views of brain connectivity (plotted using the BrainNet Viewer Toolbox^33^). In both cases, connections and nodes belonging to the same community are plotted using the same color code while grey edges correspond to connections between different modules. The size of the nodes is given by the degree.

**Figure 5.**
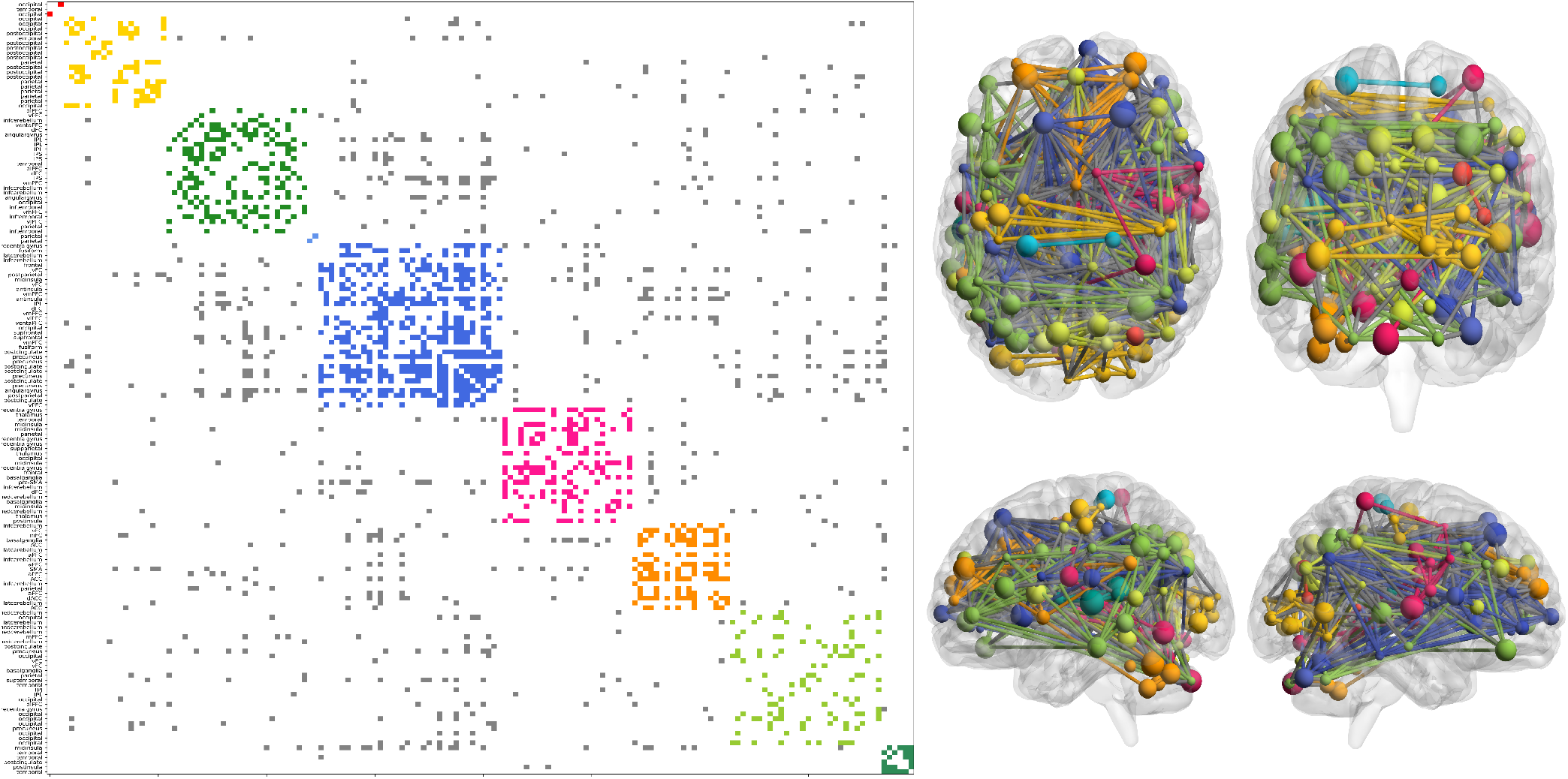
Graph partitioning of the communities defined by using QA for the Dosenbach atlas. Left: Reordered connectivity matrix according to module assignment. Right: Axial, coronal, and 2 hemispheres sagittal views of brain connectivity (plotted using the BrainNet Viewer Toolbox^33^). In both cases, connections and nodes belonging to the same community are plotted using the same color code while grey edges correspond to connections between different modules. The size of the nodes is given by the degree.

For the AAL atlas connectome, if we look for 5 communities as depicted in Fig. 4, those appear to be almost the traditional brain lobe subdivision: subcortical regions (yellow), occipital lobe (red), temporal medial (cyan), fronto-temporal medial including the default mode network (blue), fronto-temporal lateral (green). The parietal lobe was spread across the neighbouring.

For the Dosenbach parcellation, despite the difficulties of the methodology for identifying meaningful partitions, a large-scale pattern was observed, which mostly differentiated visual, sensory-motor and dorsal attention regions. More specifically, in Fig. 5 we can see depicted 9 clusters: the visual network comprising the occipital lobe but also some parietal regions (yellow) as previously described^37^, lateral temporal-parietal (mild green), extended default mode network (DMN) (dark blue), mostly thalamus and basal ganglie (magenta), cerebellum, anterior cingulate cortex and supplementary motor area (orange), superior temporal (light green), some region of post-insula and temporal (dark green). The red cluster was connecting only 2 regions in the occipital lobe, and the light blue cluster only 2 regions in the parietal lobe. The acronyms of the brain regions are the same as reported in^35^.

## 4 Discussion

The human brain exhibits an organization of a *small-world network*, with segregated modular regions integrated by some hubs nodes^11^. The use of connectomes as biomarkers has seen increasing use, and clustering features have also been introduced as both functional and structural network changes according to brain diseases^38^. Indeed, data driven approaches have being increasingly implemented to investigate relationships between brain connectivity pathology, for instance Alzheimer’s disease^39^. Therefore, it is pivotal to increase the precision of clustering brain connectivity representations.

In the presented work we investigated the capacity of a quantum annealer to perform clustering on brain connectivity data. We focused on comparing the clusters obtained with the same data by using classical and quantum computing in a D-Wave machine. The modularity metric was used to assess the quality of a community organization. This metric has values between 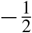 and 1, where the highest value is considered an indicator of better topological organization^2,31^.

The maximum number of clusters to detect is a parameter required by both algorithms we tested. Yet, in both cases, the algorithm was free to assign each node to any module which, in some cases, resulted in empty communities. In practice, this meant that both classical and quantum approaches could partition the graph in any number of communities only bounded by the maximum number of modules allowed. Communities obtained using the quantum device showed significantly higher modularity indices. This is in agreement with previous works comparing clustering with different datasets and similar settings^21^.

Interestingly, the number modules found by the quantum device was consistently smaller despite reaching higher modularity. We might argue that LCDA overestimated the number of clusters resulting in less edges between nodes inside a given community. Were this to occur, the modularity index would decrease, since the number of edges expected by chance, as quantified in Eq. (5), remains unaltered. Furthermore, in the two tested brain graphs, the *k*-th eigengap corresponding to the assignment of *k* clusters was higher (Fig. 3 (b) and (c)), which might be indicative of the significance of the obtained partition. Nonetheless, the lack of ground-truth for brain cases represents a limitation.

A puzzling question arises when trying to understand why QA finds higher modularity indices. To this end, we note that enforcing both QA and LCDA to find 2 communities in the Karate club graph, the results were identical. We may hypothesize that the energetic landscape to explore in these limited conditions is simple, hence both algorithms successfully find the unique isolated minimum (e.g., minimum of a parabolic function). However, the unsupervised problem (i.e., not imposing a fixed number of communities) or the presence of large networks may add significant complexity^40^. Crucially, QA does not rely on exact shape of the landscape^28^ hence providing a possible bypass to this issue (Fig. 1 (a)).

Compared to the previous studies based on the binary subdivision of hierarchical clustering^21,41^, our approach can handle odd numbers of clusters and settings where the value *k* is not a power of 2. Moreover, it is worthwhile to mention that the quantum results are stable even considering potential hardware problems known in this type of devices^42^, indeed we found a close to zero standard deviation repeating the experiments 100 times.

In our experiments with the brain connectome, we used two different types of brain connectome (coming from a structural and functional parcellation) to test our algorithm by investigating differences. Interestingly, in both cases, the quantum algorithm obtained higher modularity. In this study, we avoided investigating the clinical outcomes related to clustering either structural or functional brain connectomes as this is beyond the focus of this paper. Nevertheless, the brain communities obtained on the quantum device significantly resembled the conventional subdivision of brain structures.

Connectivity based approaches tend to generally find at least 6 minimal clusters^43^. As we also found, one cluster is generally represented by the visual system (occipital cluster). In fact, even at coarse scale (4-6 clusters), the visual system generally represented a separate community from the somatosensory system and others^44^. Somatomotor communities usually remains connected. As in our results there is a cluster spanning several midlines (medial prefrontal cortex and posterior cingulate cortex), defining the so called default mode network: A popular network involved in wakeful rest, daydreaming and mind-wandering^45^. This division was present in both the structural and functional atlas based connectome of our experiments.

A concerning limitation of the proposed approach was the computational time. We noticed that the quantum annealer took a considerably larger amount of time than its classical counterpart. The speed was not dependent of the communication between the client and the quantum computer, but we believe it was related to the current configuration of the qubits and the solver. Successfully performing QA in system of qubits relies on smooth, adiabatic transitions (i.e., quantum tunneling) between the ground states of two energetic landscapes (see Eq. (12)). Eventually, these transitions govern the computational complexity and the time^46^. Several factors can influence this complexity, including the numbers of nodes in a graph. However, solutions found through QA were more stable than expected, with not much noise and even achieving higher scores. Physical limitations on the D-Wave platforms currently in use include finite precision, sparse connectivity, and a finite amount of qubits which are connected in a Chimera graph. It can be hypothetized that having more qubits during computation and increased connectivity will improve the performance of this technique. In our approach, the community detection problem for the Zachary graph problem has been solved by using a more elegant solution than in the paper^21^. In that approach, a challenging aspect was also given splitting the graph into pieces supported by the combined effect of *qbsolv* ^2^ and the annealing process. The problem was to prepare a graph suitably sliced. This work required 300 × 4 × 4 = 4800 Chimera edges (Fig. 1 (b)). Used in this work D-Wave 2000Q can embed a fully connected 64 node graph that required at least 4096 Chimera edges. This indicates that the 300 edges thresholded version of the Zachary graph is comparable to the maximum fully connected graph that can be embedded in the D-Wave 2000Q machine. In our case, the maximum value of variables we can use is equal to 5000. Moreover, the reduction itself is significantly more straightforward, as we do not need to use graph slicing.

As a further study we will expand individual analysis of brain connectomes to multi-layer connectomes^14^ and to common eigenspaces of networks from population studies^9^.

## 5 Conclusion

In this article, we have shown the power of a quantum computer, in particular a community detection as an optimization issue to be solved by quantum annealer for brain community detection. The results analysis shows that the quantum computer is capable of rendering a highly efficient community structure with modularity superior to classical computing. One of the most noteworthy results from applying this QA process is that the community structure is obtained “all at once” in the technique within the annealing period and we can compute any number of clusters in contrast with previous approaches which could compute only power of 2 number of clusters.

## 6 Acknowledgments

This research is supported by the European Union’s Horizon 2020 research and innovation programme under grant agreement Sano no 857533, and by the International Research Agendas programme of the Foundation for Polish Science, co-financed by the European Union under the European Regional Development Fund. This research was tested on D-Wave quantum computing.

## Supplementary Material

**Table S1.** Summary of the properties of the used networks

| Network | size | number of edges | average degree |
| --- | --- | --- | --- |
| Karate club | 34 | 78 | 4.6 |
| AAL90 | 90 | 441 | 9.14 |
| Dosenbach | 160 | 891 | 11.14 |

**Figure S1.**
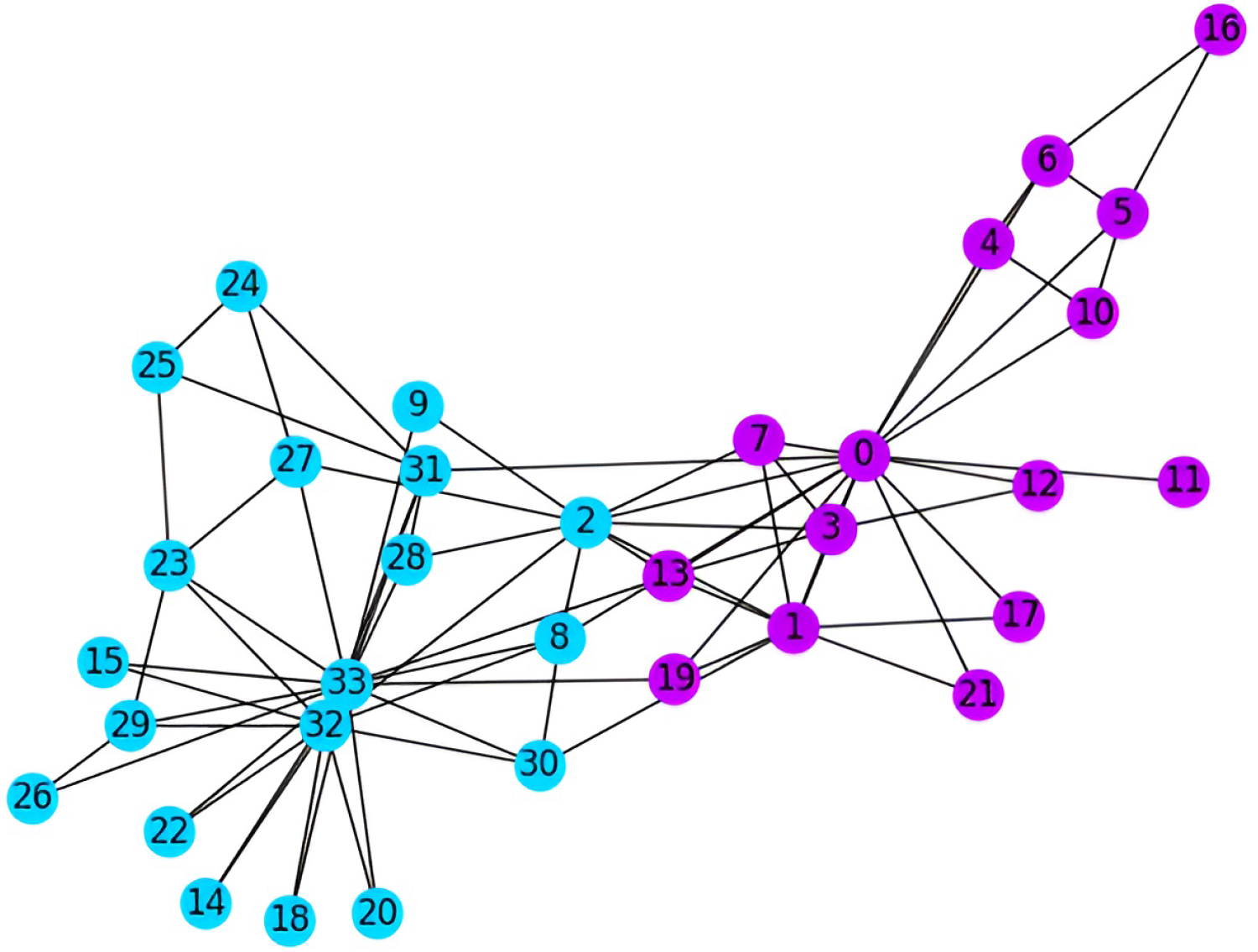
Ground-truth labeling for the communities in the Karate club^32^

## Footnotes

1 https://docs.ocean.dwavesys.com/en/stable/concepts/dqm.html

2 https://docs.ocean.dwavesys.com/projects/qbsolv/en/latest/

